# Nuclear DNA-encoded fragments of mitochondrial DNA (mtDNA) confound analysis of selection of mtDNA mutations in human primordial germ cells

**DOI:** 10.1101/2021.10.18.464832

**Authors:** Zoë Fleischmann, Sofia Annis, Melissa Franco, Sergey Oreshkov, Konstantin Popadin, Dori C. Woods, Jonathan L. Tilly, Konstantin Khrapko.

**Affiliations:** Laboratory of Aging and Infertility Research, Department of Biology, Northeastern University, Boston, MA, USA; Center for Mitochondrial Functional Genomics, Immanuel Kant Baltic Federal University, Kaliningrad, Russian Federation; Center for Integrative Genomics, University of Lausanne, Lausanne, Switzerland

## Abstract

The resilience of the mitochondrial genome to a high mutational pressure depends, in part, on purifying selection against detrimental mutations in the germline. It is crucial to understand the mechanisms of this process. Recently, Floros et al. concluded that much of the purifying selection takes place during the proliferation of primordial germ cells (PGCs) because, according to their analysis, the synonymity of mutations in late PGCs was seemingly increased compared to those in early PGCs. We re-analyzed the Floros et al. mutational data and discovered a high proportion of sequence variants that are not true mutations, but originate from NUMTs, the latter of which are segments of mitochondrial DNA (mtDNA) inserted into nuclear DNA, up to millions of years ago. This is a well-known artifact in mtDNA mutational analysis. Removal of these artifacts from the Floros et al. dataset abolishes the reported effect of purifying selection in PGCs. We therefore conclude that the mechanism of germline selection of mtDNA mutations remains open for debate, and more research is needed to fully elucidate the timing and nature of this process.

## Introduction

Mitochondrial DNA (mtDNA) is known for having a high mutational rate and a high density of crucial genes. Therefore, mtDNA mutations not only underlie a spectrum of mitochondrial diseases (Lander and Lodish, 1990), (Elliott et al., 2008) but also, if left unchecked, could result in long-term detrimental effects on species evolution (Popadin et al., 2007). It is thus vital to understand how mtDNA mutations are kept at a sustainable level through generations. A seminal study from 2008 demonstrated that significant levels of purifying selection against detrimental mutations occur in the germline (Stewart et al., 2008). Mutations are depleted of nonsynonymous variants as early as the second generation after mutations are generated *de novo* in a mtDNA ‘mutator’ mouse line. However, the mechanism(s) and precise timing of germline purifying selection remain unknown. More recently, Floros and colleagues published a study of mtDNA mutations in human embryonic primordial germ cells (PGCs), which asserted that purifying selection acted at an early stage in the developing germline during PGC proliferation and migration (Floros et al., 2018). This assertion was based primarily on an observation of a sharp increase of average synonymity of mtDNA mutations in late-stage PGCs compared to early-stage PGCs.

## Results and Discussion

### Comparison of mutations to known NUMTs reveals significant contamination with NUMT sequences

Careful review of the Floros et al. study, revealed that their key result, which was a relative increase in synonymity in late-stage PGCs, was unusual in that it appeared to result not from a depletion of nonsynonymous mutations – as would be expected under negative purifying selection- but instead from an increase in the frequency of synonymous mutations. Further examination of their primary mutational data shown in Table S5 of their supplementary information unexpectedly revealed that a majority (~80%) of sequence variants were found in many unrelated samples. Such recurrence is generally not expected of true nascent mutations, which are individually rare events and therefore have negligible probability to occur simultaneously and independently in multiple samples. We reasoned that this inter-sample repetition of low fraction sequence variants may actually be due to nuclear DNA of mitochondrial origin (NUMT) contamination, a long-known problem of mtDNA mutational analysis (Hu and Thilly, 1994), (Khrapko et al., 1994), (Hirano et al., 1997). NUMTs are nuclear pseudogenes of mtDNA, many of which are millions of years old and have accumulated many sequence differences from the true mitochondrial genome. Our alignment of Floros et al. variants to several known NUMT sequences indeed revealed at least 2 contributing NUMTs. In fact, a single one of these NUMTs located on chromosome 5 was responsible for ~20% of protein coding variants reported by Floros et al. as real mutations. We further found that NUMT contamination was most likely introduced by co-amplification of the NUMT sequences by the PCR approach used in their study (**Figure 1**). Interestingly, we have previously studied this NUMT and its homologs in great apes and demonstrated that it reveals distant interspecies hybridization event in the evolutionary history of hominids (Popadin et al., 2017).

**Figure 1.**
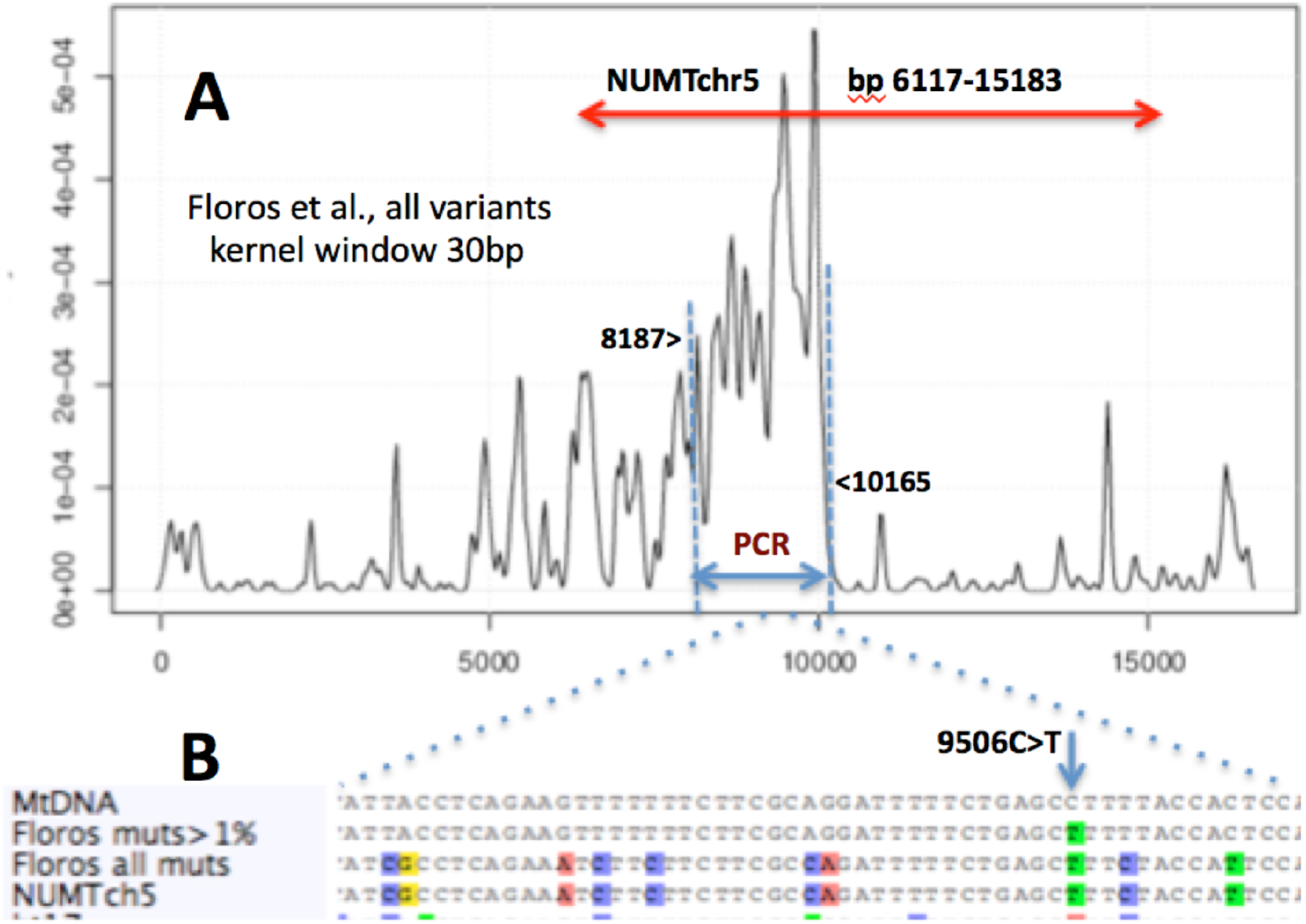
NUMT contamination in PGC mtDNA mutational dataset: spatial distribution and alignment of sequence variants detected in PGCs. **A.** Frequency distribution of Floros et al. variants along the mitochondrial genome. The unexpected dense patch of variants (between ~8kb and 10Kb, dashed vertical blue lines) corresponds to one of the PCR fragments used in Floros et al. (bp:8187-10165, horizontal blue double arrow). The PCR primers of this fragment happen to co-amplify a portion of the nuclear pseudogene of mtDNA (NUMT) on Ch5 (hg38(100045938-100055045), red double arrow). **B.** A representative section of alignment of reference mtDNA sequence with Ch5 NUMT sequence shows that a great majority of Floros et al. variants in this genome region are identical to the NUMT-derived variants, which corroborates their NUMT origin. ‘Floros all muts’: a mtDNA sequence where we incorporated **all** variants from Table S3 of Floros et al., including those that are under the 1% cutoff. ‘Floros muts >1%*’:* Table S3 variants of Floros et al. were limited to those above the 1% cutoff. Of 10 variants located in the sections of the genomes shown here, one (9506C>T, blue down arrow) exceeded the threshold and thus was erroneously considered a confirmed mutation. This illustrates the mechanism of ‘leakage’ of NUMT-driven variants into the pool of ‘confirmed mutations’.

To further corroborate the NUMT-based explanation of the Floros et al. findings, we searched the raw sequencing data of Floros et al. for typical NUMT-derived reads, which are traditionally recognized by the presence of multiple nucleotide changes within short sequence stretches (Hirano et al., 1997), (Khrapko et al., 1994). This feature reflects the fact that highly diverged ancient NUMT sequences carry a high density of sequence changes (compared to the contemporary human mitochondrial genome). For example, the NUMT at chromosome 5, which shows as a part of contamination in Floros study, contains patches of ~10 changes in 100 nucleotides. To determine if the presumably NUMT-derived sequence changes are arranged as such characteristic patches, the primary next generation sequence reads were extracted from the Floros et al. dataset. We then performed multiple alignment of the raw read sequences against the sequence of the NUMT at human chromosome 5 (See **Methods** for details). As expected, this analysis revealed a significant proportion of sequence reads characteristic of NUMTs, with patches of multiple mutations (**Figure 2**). Collectively, these results undoubtedly imply that NUMT-derived variants represent a substantial part of the Floros et al. mutation dataset.

**Figure 2.**
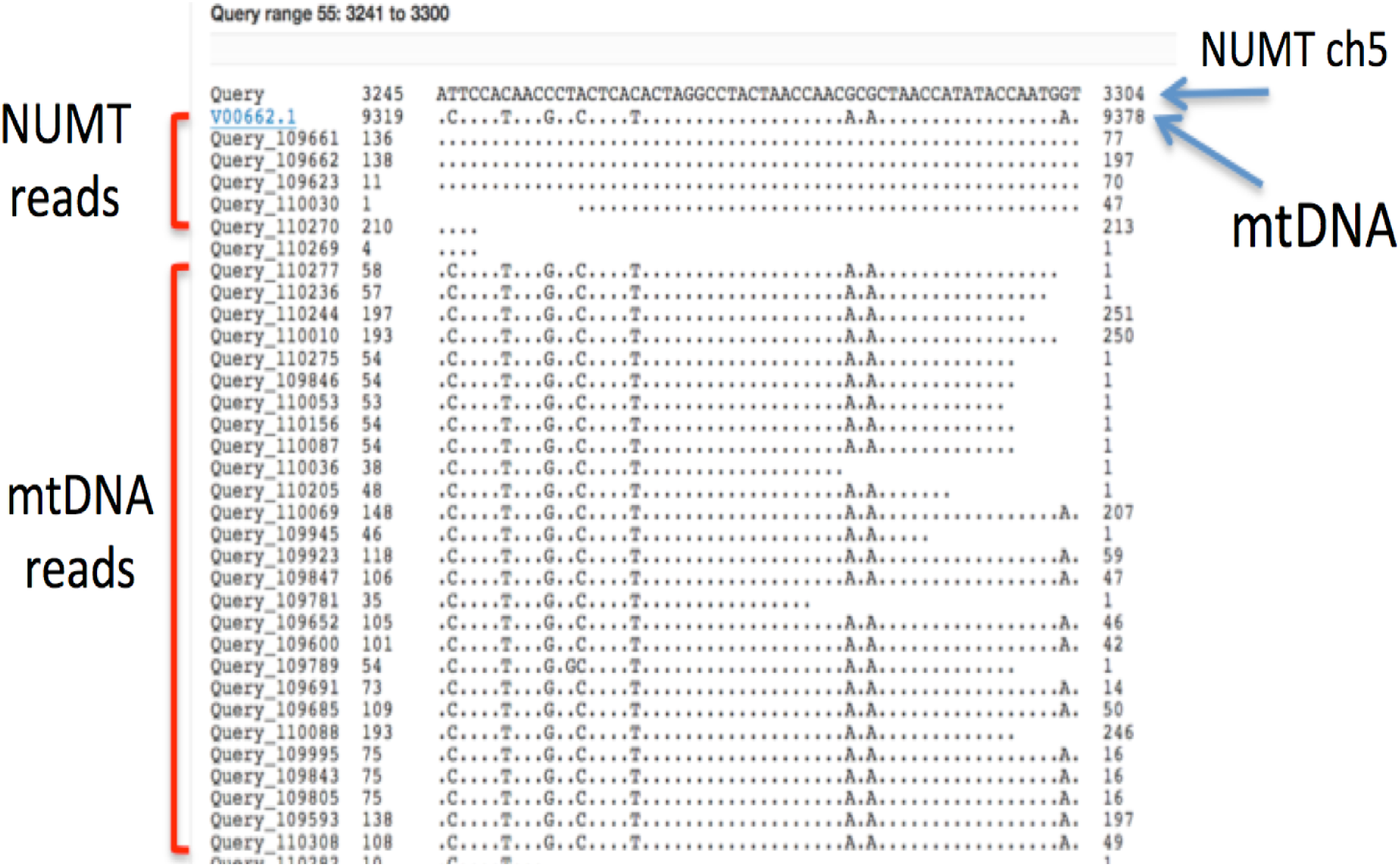
A significant proportion of primary next generation reads in the Floros et al. data are NUMT reads identified by the presence of multiple, NUMT-specific sequence variants. Shown here is a representative fragment of an alignment of the primary reads from the NUMT-rich late PGC sample from Floros et al. to the genomic sequence of the NUMT located on chromosome 5 (hg38(100045938-100055045)). In this section of the alignment, NUMT reads show up to 8 NUMT-specific sequence variants per read. Note that NUMT sequence changes appear as if these happened in mtDNA reads. This is because NUMT sequence was used as query in this BLAST alignment to keep NUMT reads on top.

There remain a few sequence variants that are repeated in several samples, for which we were not able to identify a known NUMT origin. These variants are likely derived from other, more recent, NUMTs, the presence of which in human nuclear genomes have been confirmed by the authors as well as other groups (Wei et al., 2020), (Lutz-Bonengel et al., 2021). Alternatively, variants repeated in unrelated samples could also be derived from a number of other sources, ranging from contamination by external mtDNA to ‘hotspots’ associated with instrumental error. In any case, we feel that repetitive variants must be considered errors, unless independent evidence is presented that these events are natural mutational hotspots. Even if mutations are proved natural hotspots, however, they are of limited use for germline selection analysis, because the dynamics of hotspot mutations is usually strongly affected by the process of mutagenesis itself, which confuses analysis of selection.

### Contamination with NUMT-derived sequence variants affects the estimates of the germline selection

We next explored if contamination with NUMT-derived sequence variants had significantly affected the key conclusions of Floros et al. study. By comparing mutant fractions of recurrent sequence variants in different samples, we found that NUMT-derived variants are present at systematically and significantly different frequencies. Most of these variants are below the 1% cutoff adopted by Floros et al. and thus are not included in the analysis. However, in some samples, NUMT-derived variants occasionally exceed 1% and thus were deemed ‘true’ mtDNA mutations. Importantly, a large majority of NUMT-derived variants that exceeded the 1% threshold were found in one particular late-stage PGC sample.

It may look surprising that NUMT-derived contamination has different levels in different samples, because as a nuclear sequence, the NUMTs copy number per cell is expected to be constant. However, the actual level of *NUMT contamination* depends on the nuclear-to-mtDNA ratio, and the copy number of mtDNA can significantly differ between samples of the same tissue type, as was demonstrated in the paper describing NUMT contamination in brain samples (Hirano et al., 1997). These differences are related to the variability of mtDNA levels and/or differences in DNA integrity between samples.

We note that NUMT-derived variants, which are in fact differences between mtDNA and the corresponding NUMT sequences, are expected to be (and are) highly synonymous. This inference may look surprising because NUMTs are essentially pseudogenes of mtDNA, and mutations in pseudogenes are conventionally thought of as nonsynonymous. Explanation for this surprising observation is that from the evolution perspective, a majority of NUMT variants are “reverse mtDNA mutations”. Indeed, on the evolutionary time scale, NUMT sequences, which reside in the ‘protective’ nuclear DNA environment, mutate at a much slower rate than the corresponding sequences in the far less protected mitochondrial genome. So a great majority of the differences between NUMTs and the contemporary mtDNA sequence are in fact mutations in *mtDNA* (rather than in NUMT) that were fixed in the species’ mitochondrial genome since insertion of the NUMT (which was a part of mtDNA genome contemporary for that time) into the nuclear genome. Thus, a majority of NUMT/mtDNA differences are historic mtDNA mutations, which are mostly non-adaptive and neutral. We have demonstrated this peculiar feature of NUMT mutations in humans (Popadin et al., 2017). This has been demonstrated previously in bristletails (Baldo et al., 2011).

Returning to Floros et al., because the sample most rich in NUMT-derived mutations happened to be a late-stage PGC sample, this sample brought a large number of synonymous NUMT-derived variants into the late-stage PGC mutation pool. This explains the puzzling surge of synonymous variants in late-stage PGCs. More importantly, this resulted in Floros et al. reaching an experimentally unsubstantiated conclusion of negative selection in PGCs. We then asked if removal of the erroneous repetitive variants could yield a ‘clean’ mtDNA mutational dataset that would allow for a more accurate assessment of the dynamics of true mtDNA mutations during human PGC development. Unfortunately, once these repetitive variants were removed, there remain only a few protein-coding mtDNA mutations in PGCs, which are not sufficient in number to make statistically significant inferences.

In summary, our results indicate that contamination of the Floros et al. mtDNA mutational data with NUMT sequences created bias in favor of their assertion of negative selection during PGC proliferation and migration. When these contaminating variants are removed, the data become statistically underpowered for analysis. We therefore conclude that the mechanism and timing of germline selection of mtDNA mutations requires more research to be fully elucidated.

## Methods

### Synonymity and repeatability analysis of sequence variants of mtDNA

The data pertaining to the Floros et al. study have been imported from the supplemental table 5 at https://doi.org/10.1038/s41556-017-0017-8, (Floros et al., 2018). Synonymity was analyzed using in house excel-based spreadsheets based on lookup function in excel. Identical mutations in different samples were identified using ‘countif’ functions.

### Extraction and multiple alignment of raw reads from the NGS dataset

We extracted raw read NGS data from NCBI Sequence Read Archive record SRR6288291. We used TrimGalore software to remove adapters and trim low quality reads. We filtered out 3.7% of bases from raw data due to low quality, removed 0.1% of sequences, which become shorter than 20bp. Adapters were removed from 29.4% of reads. Next, we aligned reads against two references - chr5 NUMT reference and mtDNA reference with Smith-Waterman alignment algorithm implemented in bwa utility (bwasw). For mtDNA alignment we filtered out alignment reads leaving in only reads a between positions 8750 and 10250 of mtDNA, in order to isolate region, which is homologous to NUMT at chromosome 5 (hg38(ch5100045938-100055045). For each read, we derived quality measure from CIGAR as a percentage of M symbols - we calculate this from SAM aligned files and do this procedure independently for each read in both mtDNA and NUMT alignment. Next we stratified reads out as aligned onto both references with quality measures greater than 0.9 for both NUMT and MTDNA references, NUMT -aligned reads as reads with mtDNA quality less than 0.8 and NUMT quality greater than 0.99, and mtDNA-aligned reads with mtDNA quality greater than 0.99 and NUMT quality less than 0.8. Last, we convert each set of reads from fastq into fasta format, leaving only nucleotide strings before BLAST operation.

*Multiple alignment of raw reads* was performed using the NCBI BLAST server with default parameters, with the sequence of the human NUMT at chromosome 5 (hg38(ch5100045938-100055045) as a query.

## References

Baldo, L., de Queiroz, A., Hedin, M., Hayashi, C.Y., and Gatesy, J. (2011). Nuclear-mitochondrial sequences as witnesses of past interbreeding and population diversity in the jumping bristletail Mesomachilis. Mol Biol Evol 28, 195–210.

Elliott, H.R., Samuels, D.C., Eden, J.A., Relton, C.L., and Chinnery, P.F. (2008). Pathogenic mitochondrial DNA mutations are common in the general population. Am. J. Hum. Genet. 83, 254–260.

Floros, V.I., Pyle, A., Dietmann, S., Wei, W., Tang, W.C.W., Irie, N., Payne, B., Capalbo, A., Noli, L., Coxhead, J., et al. (2018). Segregation of mitochondrial DNA heteroplasmy through a developmental genetic bottleneck in human embryos. Nature Cell Biology 2018 20:2 20, 144–151.

Hirano, M., Shtilbans, A., Mayeux, R., Davidson, M.M., DiMauro, S., Knowles, J.A., and Schon, E.A. (1997). Apparent mtDNA heteroplasmy in Alzheimer’s disease patients and in normals due to PCR amplification of nucleus-embedded mtDNA pseudogenes. Proceedings of the National Academy of Sciences 94, 14894–14899.

Hu, G., and Thilly, W.G. (1994). Evolutionary trail of the mitochondrial genome as based on human 16S rDNA pseudogenes. Gene 147, 197–204.

Khrapko, K., Andre, P. C., Cha, R., Hu, G., and Thilly, W.G. (1994). Mutational spectrometry: means and ends. Prog Nucleic Acid Res Mol Biol 49, 285–312.

Lander, E.S., and Lodish, H. (1990). Mitochondrial diseases: gene mapping and gene therapy. Cell 61, 925–926.

Lutz-Bonengel, S., Niederstätter, H., Naue, J., Koziel, R., Yang, F., Sänger, T., Huber, G., Berger, C., Pflugradt, R., Strobl, C., et al. (2021). Evidence for multi-copy Mega-NUMTs in the human genome. Nucleic Acids Research 49, 1517–1531.

Popadin, K., Gunbin, K., Peshkin, L., Annis, S., Fleischmann, Z., Kraytsberg, G., Markuzon, N., Ackermann, R.R., and Khrapko, K. (2017). Mitochondrial pseudogenes suggest repeated inter-species hybridization in hominid evolution. bioRxiv 134502.

Popadin, K., Polishchuk, L.V., Mamirova, L., Knorre, D., and Gunbin, K. (2007). Accumulation of slightly deleterious mutations in mitochondrial protein-coding genes of large versus small mammals. Proceedings of the National Academy of Sciences 104, 13390–13395.

Stewart, J.B., Freyer, C., Elson, J.L., and Larsson, N.-G. (2008). Purifying selection of mtDNA and its implications for understanding evolution and mitochondrial disease. Nat Rev Genet 9, 657–662.

Wei, W., Pagnamenta, A.T., Gleadall, N., Broxholme, J., Odhams, C.A., Ambrose, J.C., Baple, E.L., Bleda, M., Boardman-Pretty, F., Boissiere, J.M., et al. (2020). Nuclear-mitochondrial DNA segments resemble paternally inherited mitochondrial DNA in humans. Nat Commun 11, 1740.

